# Integrative genomic analysis of early neurogenesis reveals a temporal genetic program for differentiation and specification of preplate and Cajal-Retzius neurons

**DOI:** 10.1101/331900

**Authors:** Jia Li, Lei Sun, Xue-Liang Peng, Xiao-Ming Yu, Shao-Jun Qi, Zhi John Lu, Jing-Dong J. Han, Qin Shen

## Abstract

Neurogenesis in the developing neocortex begins with the generation of the preplate, which consists of early born neurons including Cajal-Retzius (CR) cells and subplate neurons. Here, utilizing the Ebf2-EGFP transgenic mouse in which EGFP initially labels the preplate neurons then persists in CR cells, we reveal the dynamic transcriptome profiles of early neurogenesis and CR cell differentiation. At E15.5 when Ebf2-EGFP+ cells are mostly CR neurons, single-cell sequencing analysis of purified Ebf2-EGFP+ cells uncovers molecular heterogeneity in CR neurons, but without apparent clustering of cells with distinct regional origins. Along a pseudotemporal trajectory these cells are classified into three different developing states, revealing genetic cascades from early generic neuronal differentiation to late fate specification during the establishment of CR neuron identity and function. Further genome-wide RNA-seq and ChIP-seq analyses at multiple early neurogenic stages have revealed the temporal gene expression dynamics of early neurogenesis and distinct histone modification patterns in early differentiating neurons. We have also identified a new set of coding genes and lncRNAs involved in early neuronal differentiation and validated with functional assays *In Vitro* and *In Vivo*. Our findings shed light on the molecular mechanisms governing the early differentiation steps during cortical development, especially CR neuron biology, and help understand the developmental basis for cortical function and diseases.

## Introduction

The mammalian cortical neurogenesis occurs on a precise time schedule during development. Neural stem cells (NSCs), the neuroepithelial cells and radial glial cells (RGCs) in the ventricular zone (VZ) first undergo symmetric cell divisions to expand, then begin to produce neurons (1, 2). The first-born neurons are generated as early as E10 in mouse and form the preplate or primordial plexiform layer, which situates above the VZ and beneath the pial surface (1, 3, 4). The preplate is then split into the marginal zone (MZ, layer 1) and the subplate by immigrating cortical plate neurons around E13.5 in mouse and the seventh to eighth week of gestation in human (5–7), setting up of the upper and lower borders of the cortical plate. The formation and splitting of preplate are the first steps in the establishment of the highly organized cortical layers.

The preplate and its derivatives play a critical role in regulating subsequent inside-out neuronal migration, cortical lamination and the establishment of topographic connections (8). Subplate neurons contribute to the guidance of corticofugal and thalamocortical axons during early development and are important for the functional maturation and plasticity of the cortical circuitry (9, 10). Cajal-Retzius (CR) neurons are the major cell type in layer 1, the most superficial cortical layer. Discovered more than a century ago by Retzius and Ramon y Cajal, CR neurons have been studied extensively (4, 11). Best known for their expression of *Reln*, CR neurons regulate the inside-out migration of cortical neurons, correct layer formation, maintenance of RGC and cortical patterning (12). Loss of *Reln* or mislocalization of CR cells contributes to defective migration and disordered cortical layer formation that are associated with human brain malformation such as lissencephaly (13) and dystroglycanopathy (14). Although functional importance of these early differentiating neurons is well recognized, characterization of cell-intrinsic, dynamic gene expression patterns in these cells remains incomplete. Recent studies have identified the genes enriched in subplate neurons by microarray and RNA-seq analysis (15–19), providing a comprehensive gene expression profile of subplate at different developmental stages, which offers valuable insight on molecular mechanisms underlying neurodevelopmental disorders (18). However, how gene expression is dynamically regulated during the transition from the preplate to CR neurons remains unclear.

We previously have shown that in the Ebf2-EGFP transgenic mice (GENSAT project) (20), the GFP signal is expressed specifically in early differentiating preplate neurons and persistent in CR cells (21). After preplate splitting, virtually all of the EGFP-positive cells exhibit morphological and molecular features of CR neurons in the embryonic and postnatal brain (21), consistent with the pattern of the *Ebf2^GFPiCre^* (22), making this line suitable for tracing the development and profiling genetic and epigenetic pattern of early born neurons including CR neurons. In addition, ablation and fate mapping studies have revealed diverse origins other than cortical neuroepithelium for CR neurons, such as cortical hem, septum, ventral pallium, which could be recognized by specific molecular markers (23–26). However, whether CR cells from different origins maintain their distinct regional identities after they reach the layer 1 in the neocortex remains to be addressed.

Single-cell gene expression profiling allows further dissection of the heterogeneity and developmental trajectory within a cell population (27, 28). While multiple types of neurons in adult cortex and hippocampus have been identified by single-cell RNA-sequencing (28, 29), a full characterization of early differentiating neurons at single-cell level has not been done before. In this study, we identified novel subpopulations of CR neurons from E15.5 neocortex using single-cell sequencing and uncovered distinct developmental states within the pure population. Analysis of CR neuron lineage revealed molecular cascades along generic neuronal differentiation to CR cell-specific fate determination. Furthermore, we used FACS-purified Ebf2-EGFP-expressing cells to probe the dynamic molecular mechanisms of preplate and CR neurons during the early stages of forebrain development. Whole genome transcriptome analysis identified the initial differences between the preplate and progenitor cells at E11.5, reflecting the genetic choice during the first cell fate decision in neural differentiation. We also recognized clusters of genes that displayed temporal changes corresponding to different developmental trajectories. Genome-wide active (H3K27ac) and repressive (H3K27me3) histone modifications in preplate neurons, CR neurons and NPCs revealed that occupancies of histone marks around TSS and distal regions were distinctly different between different cell types, and unique histone modification pattern emphasized on promoter regions to reinforce CR neuron specification, suggesting epigenetic role in acquisition and maintenance of cell type-specific identities. By bioinformatic functional enrichment analysis, we also identified and validated the function of coding genes and lncRNAs that were enriched in CR neurons during neuronal differentiation. Hence, our study identify the three distinct development states within pure CR population and uncover molecular hallmarks along CR neuron differentiation, which has never been reported before. Besides, we discover unique histone modification pattern which particularly emphasize on promoter region reinforcing CR neuron specification. Accompany with them, we unravels the remarkable temporal dynamics in gene expression during differentiation of a specific neuronal subtype and provides molecular markers to extend the repertoire of preplate and CR neuron markers. Our findings will help understand the etiology of diseases such as Lissencephaly, Alzheimer’s disease, and Schizophrenia.

## Results

### Single-cell analysis identify heterogeneities within pure CR neuron population

We have previously shown that in the Ebf2-EGFP transgenic mice, EGFP is specifically expressed in layer 1 neurons and more than 95% Reln+ cells are Ebf2-EGFP+ at E15.5 by immunostaining of cerebral cortex sections (21). The Ebf2-expressing cells located in the layer 1 of the embryonic cerebral cortex consist of CR neurons from different origins including cortical hem, septum and ventral pallium (22, 25, 30). To investigate the molecular heterogeneity and characterize the developmental states of CR neurons at single cell level, we performed single-cell RNA-seq (10X genomic Chromium) of 3000 Ebf2-EGFP+ cells purified by FACS of E15.5 cortical cells. After filtering out potential noises as multiplet cells and mitochondria genes, then normalization and PCA analysis, the acquired gene expression data sets were selected for the following analysis (Fig S1A-C). By K-Means clustering, Ebf2-EGFP positive cells at this stage were molecularly clustered into eight major subpopulations, well separated and visualized in the t-SNE plot (Fig 1A and B). These eight cell clusters had distinct gene expression patterns and were distinguished with highly differential expressed markers (Fig 1A; Fig S1D). Notably, Cluster 2 was the most segregated group from the others, with high expression level of typical CR markers such as *Reln* (Fig 1A). Interestingly, we found that 46% of E13.5 CR genes and 50.6% of P2 CR genes identified from a previous study (25) were included in Cluster 2 (29 genes, and 44 genes, respectively), indicating high enrichments of common CR genes in this cluster.

**Figure 1.**
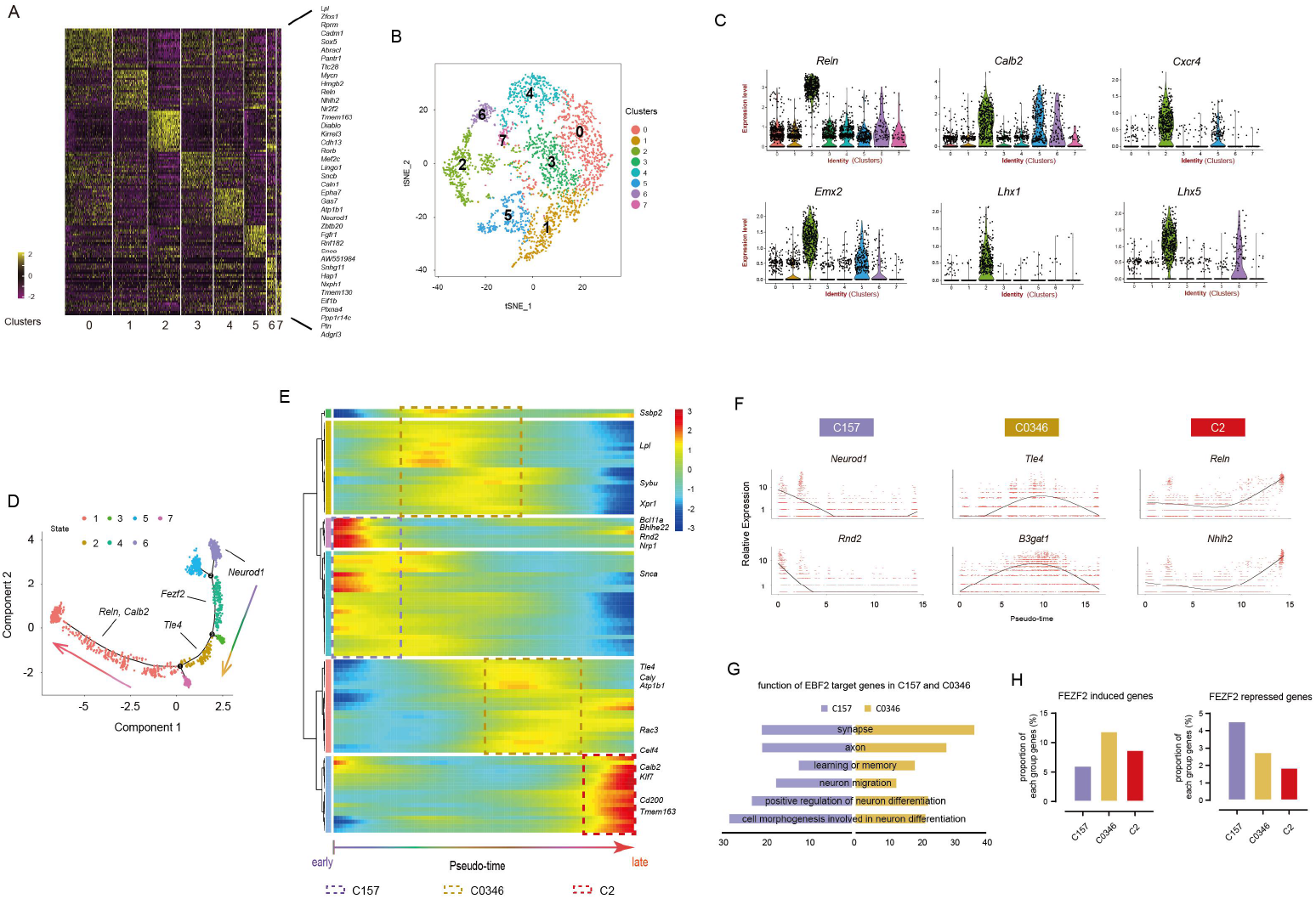
Single cell RNA-seq identify heterogeneities within CR neuron population and reveal genetic program for early neuron differentiation. (A) Heatmap of differential expressed genes in E15.5 Ebf2-EGFP+ cells sub-clusters. (B) The K-means algorithm-based plot showing eight major populations. (C) Violin plots showing genes expressed in CR neuron origins with high and specific expression in some cell sub-clusters. Each dot represents a single cell. (D) Pseudotime trajectories of Ebf2-EGFP+ cells computing reconstructed by Monocle. Cell states are indicated by colors and sequential numbers in the black circles. (E) Heatmap shows gene dynamics along pseudotemporal cortical neuron differentiation. X-axis shows pseudotime ordering from early to late, Y-axis shows a single gene expression level dynamic per row. Gene expression level from high to low is indicated by colors from red to blue. (F) Expression level dynamics across pseudotime ordering of C157 genes *Neurod1*, *Rnd2*, C0346 genes *Tle4*, *B3gat1*, and C2 genes *Reln*, *Nhlh2* are shown. (G) Functional GO analysis for EBF2 targeting genes in C157 and C0346, respectively.(H) The distribution of FEZF2 targeting induced genes and repressed genes across C157, C0346 and C2.

CR neurons were reported to be generated from many areas of developing pallium and were redistributed on the cortical surface via tangential migration (23, 25, 26, 31). It remains elusive whether CR neurons arising from different origins are molecularly distinguishable and maintain their original identities or develop to the same type of cells with identical gene expression profile after they reach the cerebral cortex. We investigated whether the eight clusters expressed different regional genes. We found that typical CR marker genes such as *Reln* and *Calb2* were expressed in all cell subpouplations. Surprisingly, we did not detect segregation of regional specific genes in the DEGs of the eight clusters. In contrast, we observed genes expressed in various CR cell original territories were enriched in the same cluster. For example, *Cxcr4* (cortical hem), *Emx2* (VZ), *Lhx1* (septum, ventral pallium), *Lhx5* (cortical hem, septum and ventral pallium) (26, 31, 32) were all included in Cluster 2 (Fig 1C; Fig S1E), suggesting single cells expressed multiple genes of different positional origins. Therefore, our results showed that different CR neuron origins can not be segregated by unsupervised molecular clustering, suggesting that CR neurons may lose the original region identity once they migrate into the cerebral cortex layer 1 and molecular and functional maturation of CR neurons may follow a general path regardless of their origins.

### Reconstruction of CR neuron developmental lineage reveal a transition from activation of a common neuronal differentiation program to repression of other neuronal subtypes to establish the CR cell-specific identity

To characterize the developmental insights of the CR neurons, we performed pseudotime reconstruction using Monocle analysis (33). We chose genes that were expressed across more than 10 cells and their average expression levels were more than 1 for this analysis (Fig 1D; Fig S1F). We found that the eight major cell subpopulations clustered from t-SNE plot corresponded to the temporal cortical neuron development progression from early to late stages (Fig 1E). Genes in Cluster 5 could be the first activated population during the pseudo-temporal ordering, sequentially followed by Cluster 1 and 7 genes, which displayed high expression levels at the beginning of the time period, later dropped to low levels. For example, Cluster 5 gene *Neurod1* is known to be expressed in committed neuronal progenitor cells at the upper SVZ and in newborn neurons at the lower IZ, which plays a prominent role in cell fate specification (34, 35). Cluster 1 gene *Rnd2* functions in the downstream of *Neurog2* to promote neuron migration (36). Other genes in these three clusters such as *Plxna4* is also reported to regulate neuronal migration in neocortex development (Fig 1E and F) (37). Thus, Cluster 1, 5, 7 (C157 in short) genes might be involved in initial transition from progenitor cells to neuronal differentiation. In contrast, genes in Cluster 2 were expressed at lower expression levels at the starting site, but higher expression levels at the end. The top differentially expressed genes in this cluster were CR neuron markers, such as *Reln*, *Trp73, Calb2, Ebf2, Lhx5*, implying a more committed CR neuron state (Fig 1E and F; Fig S1G). Cluster 0, 3, 4, 6 (C0346) genes were in an intermediate state, displaying higher expression levels during the middle period of the pseudo-temporal axis than the starting and terminal points. Other layer-specific neuron markers fell into C0346, showing transient expression in Ebf2-EGFP+ cells. For example, the deep layer neuron marker *Fezf2* and *Tle4* were enriched in Cluster 3 and 4 respectively (Fig 1E and F; Fig S1G). Therefore, our pseudotiming analysis imply that differentiation of CR neurons starts first from a general neuronal differentiation program, then goes through a refinement of gene expression to suppress other cell type-specific genes, and finally reaches an establishment of the CR-specific genes.

### Transcriptional factors involved in transitions in gene expression during constraining terminal cell fates to CR neurons

To understand the molecular mechanisms of general neuronal fate committed to CR neurons identities, we focus on the genetic network of some key transcription factors that may predominate the differentiation progression. *Ebf2* is enriched in Cluster 2 and its target genes has been identified in Neurog2-induced neurogenesis from mESC assays recently (38). We found that 457 of the early phase C157 genes and 611 of the intermediate C0346 genes were the target genes of EBF2 during neural differentiation, implying that it might affect the molecular pathways of CR neuron differentiation (Table S1). Through GO analysis, we found the target genes in C157 were involved in neuron differentiation and neuron migration, while the target genes in C0346 were more correlated with synapse, axon, learning and memory properties (Fig 1G). Similarly, 132 genes in C157 and 172 genes in C0346 were CR marker ZIC2’s (Cluster 2) target genes identified from a neurogenesis assay (Fig S1H; Table S1) (39). Therefore, it raised a possibility that these genes might interact and regulate non-CR genes to allow CR-specific traits to be established. On the other hand, *Foxg1*, a gene known to suppress CR cell fate (40, 41), highly expressed in C157 and C0346 but low in C2 (Fig S1I). In our results and former studies, knockdown *Foxg1* also significantly increased C2 genes expression like *Reln*, *Ebf2*, *Nhlh2* and promoted CR neuron generation *in vitro* and *in vivo* (Fig S2F) (40–42), while *Ebf2* suppressed the expression level of *Foxg1* (Fig S2E). This implies suppression of C157 and C0346 genes facilitate C2 genes. Interestingly, *Fezf2*, an intermediate state C0346 gene, is known to encode a key transcription factor for deep layer neurons, regulating the expression of gene sets that were responsible for establishing mouse corticospinal motor neurons identities (43). Here, we found that FEZF2 induced genes were more frequently appearing in C2 than C157, while FEZF2 repressed genes were less in C2 than in C157 (Fig 1H; Table S1), suggesting that the C0346 gene FEZF2 might induce CR neuron identities via activation of C2 genes and inhibition of the general cortical neuron fate by repressing C157 genes. Taken together, a subset of key transcription factors might play a pivotal role in controlling distinct differentiation events through combinatorial actions on a variety of temporal genetic targets and networks, thus guiding initial generic cortical neuron program to terminal CR neuron-specific cell fate determination.

### Characterization of Ebf2-EGFP-expressing cells in the early cerebral cortex

To further investigate the molecular mechanisms and temporal dynamic gene network during the early stages of neural differentiation, we performed bulk RNA-seq of Ebf2-EGFP+ and Ebf2-EGFP- cells at multiple early neurogenic stages as E11.5, E13.5 and E15.5. We previously have shown that at E11.5, Ebf2-EGFP+ cells are mostly preplate neurons including both subplate neurons and CR neurons, and at E13.5, Ebf2-EGFP expression is diminishing in subplate neurons but maintain in CR neurons. By E15.5, Ebf2-EGFP+ cells are mostly CR neurons (21). Here we purified Ebf2-EGFP-expressing cells by FACS from the embryonic mouse forebrain at E11.5, E13.5 and E15.5, to compare the developmental gene expression dynamics during preplate formation and differentiation of early neurons (Fig 2A). At E11.5, Ebf2-expressing cells consisted of 17.10% ± 1.07% of the total cells. At E13.5 and E15.5, the percentage of Ebf2-expressing cells gradually decreased to 8.37 ± 2.29% and 3.90 ± 1.59%, respectively (Fig 2B). 100% of FACS-sorted Ebf2-EGFP+ cells were β-III tubulin+ at all stages, indicating these cells have committed neuronal fate decision, consistent with our previous in vivo analysis (21). Accordingly, after culturing sorted and unsorted cells for 4 days, almost all of the Ebf2-EGFP+ cells remained single and extended processes, while the Ebf2-EGFP- cells proliferated and generated large clones similar to the unsorted cell group, indicating a good separation of differentiating neurons from progenitor cells (Fig S2A).

**Figure 2.**
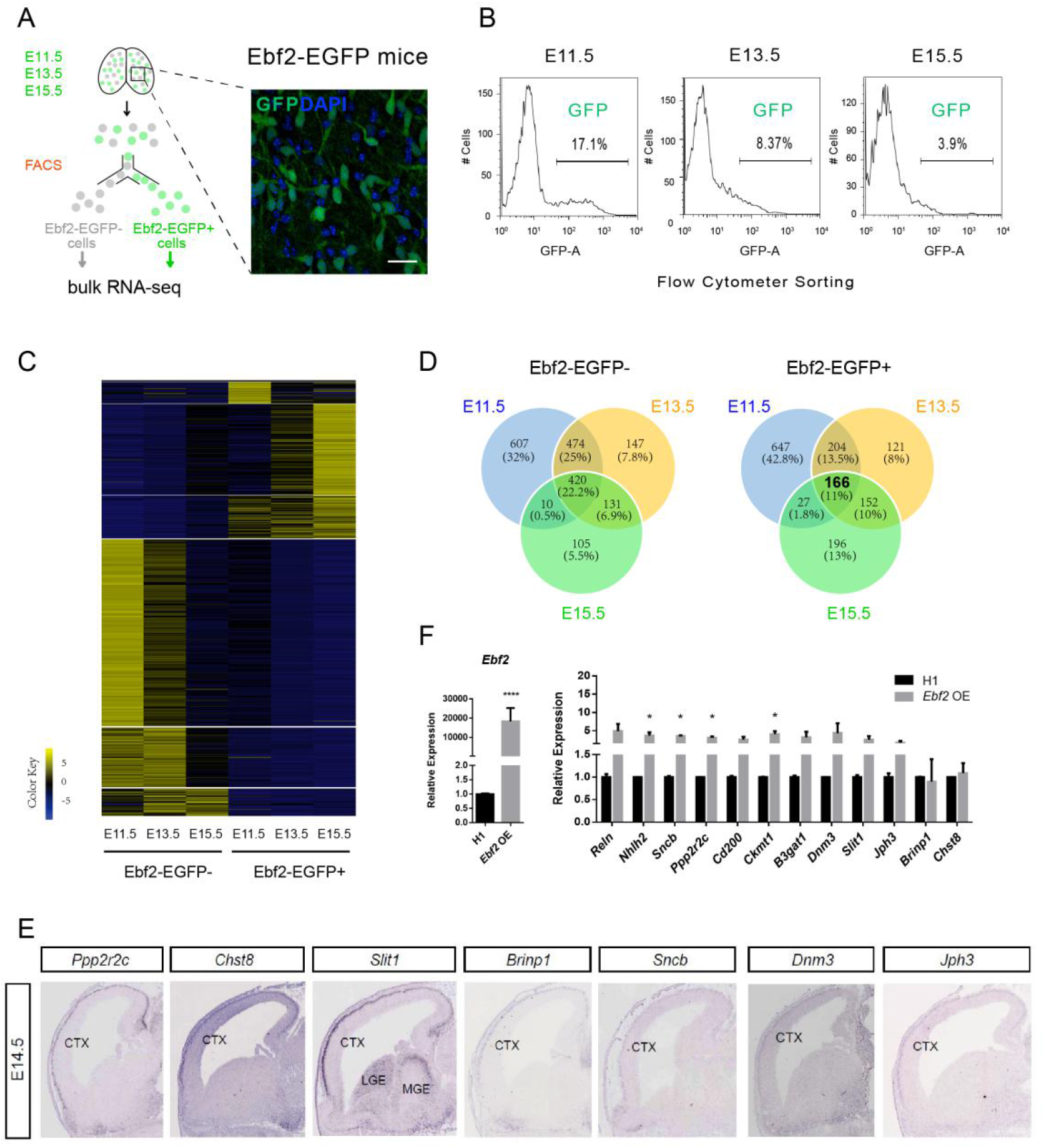
Transcriptome sequencing of Ebf2-EGFP+ and Ebf2-EGFP- cells in the early embryonic stage. (A) Schematic illustration of FACS purification of Ebf2-EGFP+ and Ebf2-EGFP- cells from early embryonic stages E11.5, E13.5 and E15.5, respectively. (B) The percentage of Ebf2-EGFP+ cells collected from FACS at each stage are calculated and shown in the graph. (C) Heatmap of all differential expressed genes (FDR<0.05) in Ebf2-EGFP+ & Ebf2-EGFP-samples at the three early embryonic developmental stages are shown. (D) Venn diagrams displaying the extent of overlapping between DEGs at the three early embryonic stages. (E) Expression of 7 CR-specific coding genes are found to be selectively expressed in cortical layer 1 of E14.5 mouse embryos (In situ hybridization images from GenePaint.org). Another CR-specific coding gene *Nhlh2* is shown in the Fig S1D. (F) qRT-PCR analysis comparing the expression levels of CR-specific genes in NSC culture assay after virus infection with control (H1) or *Ebf2*-overexpression. Data represent mean ± SEM (n=3 independent experiments, *P<0.05, Student’s T test).

### Identification of Ebf2-EGFP+ cell-enriched genes in the developing mouse forebrain

Next, we collected FACS-selected Ebf2-EGFP+ cells and Ebf2-EGFP- cells from the three early neurogenic stages and isolated total RNA for whole-genome transcriptome sequencing. Two biological replicates for each cell type (Ebf2-EGFP+ and Ebf2-EGFP-) were sequenced and expressed transcripts were identified as described in Methods. To verify the reproducibility of our samples, we analyzed the Pearson’s correlation coefficient (Pearson’s r) for the replicates and there was a strong association between different preparations of FACS-sorted cells from the same stage (Fig S2B).

To validate the RNA-seq results, we selected several coding genes, including CR neuron markers (*Ebf2* and *Calb2*) and genes associated with CR neurons fate regulation (*Foxg1*, *Tbr1*, *Ebf1*, and *Ebf3*) (11, 21, 25, 41, 42, 44), and verified by qPCR that the temporal dynamic expression levels of these genes were in accordance with that detected by RNA-seq (Fig S2C). Furthermore, the temporal expression trends were consistent with the mRNA expression pattern revealed by in situ hybridization (ISH) from the data available in Allen Brain Atlas (Fig S2C).

We used DE-Seq to analyze the dynamic expression profiles during the early cortical differentiation period. 4859 genes exhibited significant difference in expression level between the two cell populations during the time course and were selected as differential expressed genes (DEGs) for further analysis (Fig 2C; Table S2).

We analyzed DEGs expressed in any of the developmental stages sequenced (E11.5, E13.5 and E15.5). Genes with expression level (FPKM) more than 1, and 2-fold higher in Ebf2-EGFP+ population than that in Ebf2-EGFP-population were defined as Ebf2-EGFP+ cells enriched genes, CR-enriched genes in short. We found that there were 1044 genes at E11.5, 643 genes at E13.5, and 541 at E15.5 enriched in Ebf2-EGFP+ cells, respectively. Considerable overlap was found between the different stages (Fig 2D). In total, 1513 genes displayed enriched expression in Ebf2-EGFP+ population at least at one stage, and 1894 such genes in Ebf2-EGFP-population. Of these, 166 genes were consistently enriched in Ebf2-EGFP+ population throughout all three embryonic stages, and 420 genes were consistently enriched in Ebf2-EGFP-population (Fig 2D). The 166 DEGs included known CR neuron markers such as *Reln* and *Trp73*, and genes known to be expressed in the CR neurons such as *Calb2*, *Tbr1*, *Ebf3*, *Cdkn1a* (11, 25, 44, 45). We further defined the DEGs with FPKM ≥ 5 in Ebf2-EGFP+ cells, and fold change ≥ 2 at all three stages as CR-specific genes. After excluding 10 subplate genes reported in previous studies (15, 17) and 14 CR genes reported in previous studies (25), 77 are novel genes with no reported association with CR neurons. Among the 77 CR-specific genes, 56 have valid ISH from public resources. We also noticed that there were genes showing high expression levels in both Ebf2-EGFP+ cells and Ebf2-EGFP- cells although their expression levels meet the fold change criteria. To obtain the genes specifically expressed in Ebf2-EGFP+ cells but not in Ebf2-EGFP- cells, we added a cutoff of FPKM ≤ 15 in Ebf2-EGFP- cells considering that CR-specific genes should be expressed at low level in Ebf2-EGFP- cells at these stages. 33 genes remained after the filtering and we confirmed that 8 with specific expression in layer 1 cells: the 8 genes are *Nhlh2*, *Ppp2r2c*, *Chst8*, *Slit1*, *Brinp1*, *Sncb*, *Dnm3*, *Jph3*, belonging to transcription factors, synapse molecules, and axon guidance molecules (Fig 2E; Fig S2D and E; Table S2). Hence, these genes could serve as CR-specific markers for future study.

Remarkably, the expressions of these 8 genes were regulated by overexpression or knockdown of *Ebf2*, and knockdown of *Foxg1* in E11.5 cortical cell culture (Fig 2F; Fig S2E and F). For example, *Sncb, Ckmt1* were significantly upregulated or inhibited by overexpression and knockdown of *Ebf2*, respectively. While *Nhlh2, Ppp2r2c* responded to *Ebf2* overexpression but not knockdown, the expression of *Slit1* and *B3gat1* were decreased by *Ebf2* knockdown, suggesting they might also be regulated by molecules involved in developmental pathways other than *Ebf2* (Fig 2F; Fig S2E and F).

Cortical CR neurons are considered to have multiple extra-cortex origins (46). We and others have shown previously that *Foxg1* suppressed CR neuron fate (41, 42), therefore we hypothesized that CR-specific genes identified by our RNA-seq analysis could be regulated by *Foxg1* expression. Indeed, silencing *Foxg1* using shRNA up-regulated the mRNA levels of *Ebf2, Reln,* and CR-specific genes (*Nhlh2, Ppp2r2c* and *Cd200*) (Fig S2F).

We performed Gene Ontology analysis using the DAVID program to predict the potential functions of DEGs that were enriched in the Ebf2-EGFP+ and Ebf2-EGFP-cell population (Fig 3A-D; Table S3). The GO terms associated with neuronal differentiation and functions were highly enriched in DEGs expressed in the Ebf2-EGFP+ cell population, including “synapse”, “neuron projection”, “neuron differentiation”, “cell morphogenesis involved in neuron”, “axonogenesis” and “neuron migration” (Fig 3A). In contrast, the GO terms enriched in the Ebf2-EGFP-cell population were mostly correlated with progenitor cell properties, such as “forebrain development”, “regulation of cell proliferation”, “growth factor binding”, “regulation of neurogenesis” and “dorsal/ventral pattern formation” (Fig 3A).

**Figure 3.**
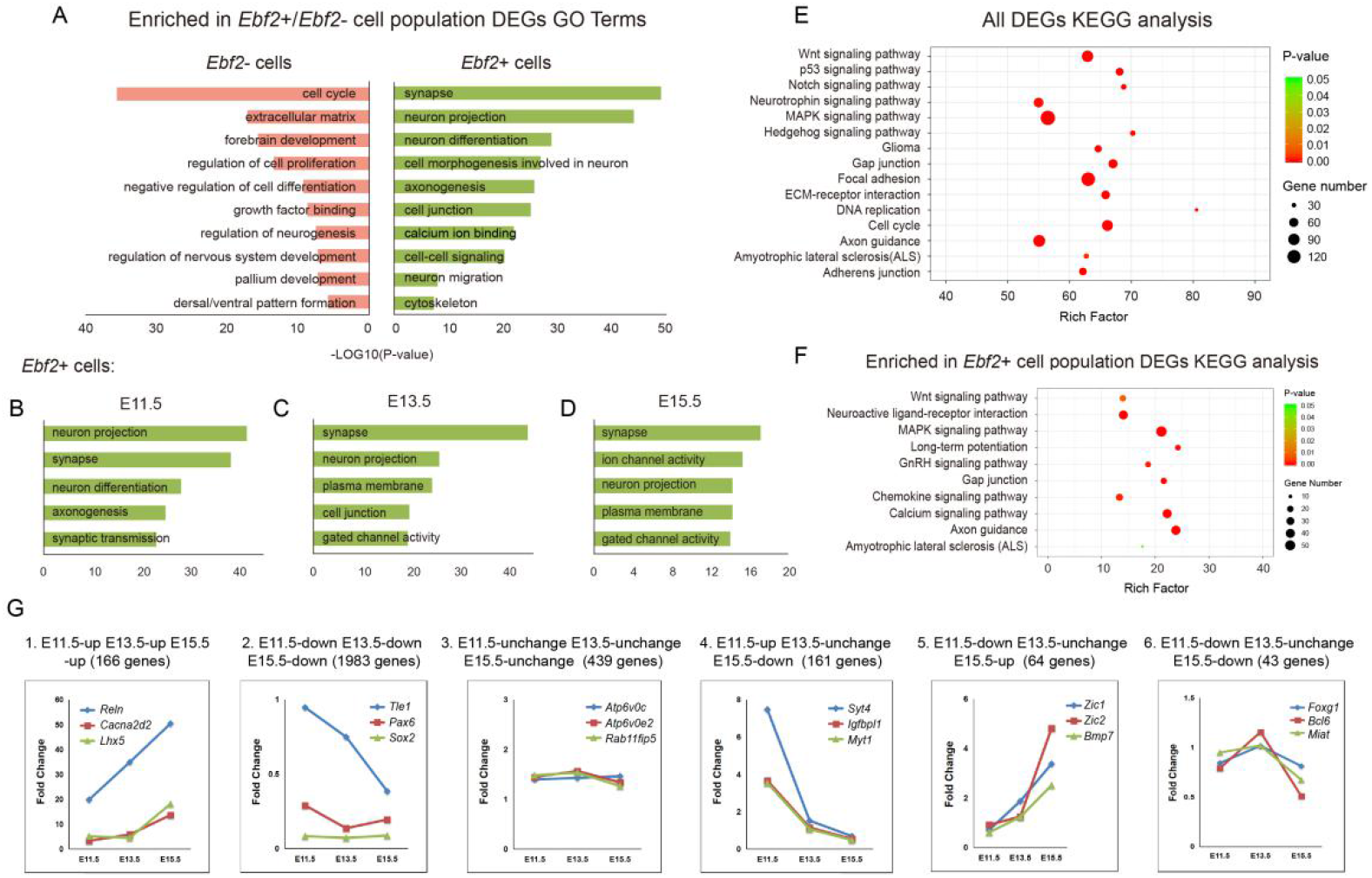
Temporal dynamic DEGs in Ebf2-EGFP+ cells and Ebf2-EGFP- cells show early neuron differentiation progression. (A) Enriched functional assessments (including biological processes, cellular components and molecular function) of highly expressed DEGs in Ebf2-EGFP+ or Ebf2-EGFP-cell population (fold change ≥ 2). (B-D) Enriched functional assessments of highly expressed DEGs in E11.5, E13.5 or E15.5 Ebf2-EGFP+ cell population, respectively. (E-F) Significantly (p<0.001, FDR<0.001) enriched KEGG pathways within DEGs from all RNA-seq samples and DEGs enriched in Ebf2-EGFP+ samples. (G) Temporal dynamic expression pattern of DEGs at the three different stages (y axis = Fold Change between Ebf2-EGFP+ cells and Ebf2-EGFP- cells). Noted the dynamic pattern is related to functions of associated genes.

In addition, we utilized the KEGG pathway repository to help determine the relative representation of specific pathways in our global RNA-seq expression data. Significantly enriched KEGG pathways of the global DEGs pointed out many signaling pathways implicated in normal neural development (Fig 3E; Table S3). For example, genes falling under the MAPK Signaling Pathway, Wnt Signaling Pathway and DNA replication were enriched. The Focal Adhesion and Axon Guidance pathways were also enriched, indicating regulation of cell morphology and neuronal differentiation were involved. Moreover, we performed the KEGG pathway analysis focused on DEGs enriched in Ebf2-EGFP+ cell population (Fig 3F; Table S3). As expected, the major groups of KEGG categories enriched in this data set were Calcium Signaling Pathway, Axon Guidance, Neuroactive Ligand-receptor Interaction, Gap Junction, GnRH Signaling Pathway, signifying differentiation of neuronal properties. Interestingly, as *Ebf2* is an important factor in GnRH neuron differentiation (47), the enrichment of GnRH Signaling pathway suggests that it would regulate similar pathway genes in cortical neurons.

### Temporal transcriptome analysis of Ebf2-EGFP+ cells reveal early neuronal specification and CR neuron differentiation dynamics

The high enrichments of these GO analysis terms and KEGG pathways imply that the transcriptional difference between the two cell populations is consistent with their different cell type identities. The components of Ebf2-EGFP+ and Ebf2-EGFP-cell population change as development proceeds. Ebf2-EGFP+ cells are mainly preplate neurons including both subplate and CR neurons at E11.5, while Ebf2-EGFP+ cells are purely CR neurons by E15.5 (21). In contrast, at E11.5 most if not all of the Ebf2-EGFP- cells are progenitor cells while at E13.5 and E15.5 the Ebf2-EGFP-population also includes cortical plate neurons. Therefore, the genes that are highly enriched in Ebf2-EGFP+ cells at all three stages, particularly those with increased enrichment, could be specifically involved in CR neuron differentiation, while the genes that are highly enriched in Ebf2-EGFP+ cells at E11.5 but not at E13.5 or E15.5 could be involved in regulating general neuronal differentiation processes. Hence, we further compared temporal changes in global DEGs at the three different stages. Genes with expression level (FPKM) in Ebf2-EGFP+ population 2 folds of that in Ebf2-EGFP-population were defined as “up-regulated”. If the ratio of FPKM (Ebf2-EGFP+/ Ebf2-EGFP-) was between 2 and 1, these genes were defined as “unchanged”. If the ratio was between 1 and 0, these genes were defined as “down-regulated”. Theoretically, we expected 27 possible combination of expression change patterns over the three time points, while our analysis showed 26 classes, missing the pattern of “E11up - E13down - E15up”. These 26 classes could be classified into 6 groups representing various aspects of early neurogenesis, and 6 typical classes were shown in Figure 3G and Figure S3 (Table S4). We hypothesized that CR-specific genes were consistently higher at the three stages, while the difference in genes involved in global neuronal differentiation would no longer be evident, after E13.5 as the Ebf2-EGFP negative population at this stage also includes newly generated cortical plate neurons. Indeed, as our expectation, the genes in the group “E11up - E13up - E15up” include the well-known CR marker genes, such as *Reln*, *Ebf2*, *Trp73* and *Lhx5*, and mature neuron genes related to axon regeneration like *Cacna2d2* (48). In contrast, genes in the “E11down - E13down - E15down” keep higher expression in Ebf2-EGFP- cells, showing that they may function on regulating NSC proliferation, such as *Sox2*, *Pax6* and *Hes5*. Genes that are first expressed in progenitor cells then in cortical neurons other than preplate and CR neurons may also be included in this group, such as *Tle1* (49). The “E11up - E13unchange - E15down” group includes subplate genes identified previously, such as *Syt4*, *Igfbpl1*, *Myt1* (15, 17), reflecting that subplate genes are highly expressed in the preplate on E11.5, and decreased in layer1 when the preplate splits later on E13.5 and E15.5. Besides, it also contains genes responsible for early neuron differentiation via radial migration like *Rnd2*, *Plxna4*. Interestingly, we found that most genes in the “E11down - E13unchange - E15up” group initially have higher expression in the cortical hem or MGE at E11.5, then appear in layer 1 at E13.5 and E15.5, implying that these genes may be involved in CR neurons migration from various origins. For example, *Zic1* and *Zic2* are expressed in the progenitor cells in the septum and cortical hem, and *Bmp7* is expressed in the cortical hem, meninges, and choroid plexus, typical CR neuron origins and location (50, 51). The genes in the “E11down - E13unchange - E15down” group include *Foxg1*, *Bcl6* and *Miat*, which are expressed along the lineage of neural progenitor cells (52, 53). The “E11unchange - E13unchange - E15unchange” group is enriched with genes associated with basic cellular organelles or activities that required for various eukaryotic cells. For example, *Atp6v0c* and *Atp6v0e2* in this group are vacuolar ATPase and play essential roles in maintaining PH in the cellular compartments, protein degradation (54, 55), *Rab11fip5* is reported to participate in protein trafficking between different organelle membranes (56). Taken together, our analysis reveal that temporal gene expression dynamics reflects distinct developmental processes, allowing us to identify novel factors that play specific roles in early cortical neurogenesis and CR development.

We further compared the temporal dynamic gene categories with three pseudo-reconstructed subpopulations (C157, C0346, C2) (Fig 1E and F) to investigate whether they are consistent in developmental dynamics along embryonic cortical neuron specification. To better represent each cluster’s specific gene expression pattern, we focused on the top 20% highly expressed genes of each cell cluster. Interestingly, we found that many “up-up-up” class genes were C2 genes, including reported CR genes (e.g. *Reln*, *Trp73*, *Lhx5*, *Calb2*, *Ebf2*, *Ebf3*, *Lhx1*, *Cdkn1a*), CR-specific genes identified in this study (e.g. *Nhlh2*, *Ppp2r2c*), and other genes related to mature neurons (*Kcnip2*, *Nxph3*, *Tmem163*) (Fig 1E and F; Fig S1G; Table S2; Table S4). Many “up-unchange-down” and “up-down-down” classes genes appeared in the early state C157 cluster, such as *Neurod1*, *Rnd2*, *Plxna4* (Fig 1E and F; Table S4), in line with their trends with higher expression at the initial stage and lower expression later. Genes in the “unchange-unchange-up” and “unchange-up-up” classes genes tended to be in the intermediate state C0346, such as *Tle4*, *Fezf2* (Fig 1E and F; Fig S1G; Table S4). Hence, both transcriptome analysis at single and bulk cell level identify a transcriptional cascade from general neuronal differentiation to specification of a neuronal subtype.

### LncRNAs enriched in Ebf2-EGFP+ cells show functional potentials for early differentiating neuron

From our RNA-seq analysis, we identified 2417 long non-coding RNA transcripts in cortical cells using the Cufflinks method. 200 lncRNAs were differentially expressed between the Ebf2-EGFP+ and the Ebf2-EGFP-population and termed DElncRNAs (Fig 4A). 58 lncRNAs were highly expressed in Ebf2-EGFP+ cells compared to Ebf2-EGFP- cells, while 64 lncRNAs showed the opposite pattern. We further filtered DElncRNAs with these criteria: 1) specifically expressed in Ebf2-EGFP+ cells with FPKM at least 2-fold of that in Ebf2-EGFP- cells (Fig 4B); these lncRNAs also showed temporal specific expression patterns. 2) lncRNAs from 1) with FPKM ≤ 15 in Ebf2-EGFP- cells during all three stages, and FPKM ≥ 5 in Ebf2-EGFP+ cells at E15.5. We identified 9 lncRNAs that met the criteria and these had never been studied before in the context of neural development (Table S5). We defined these lncRNAs as CR-specific lncRNAs and called *Ln-CR*# for simplicity. The qPCR results to validate the mRNA expression level were generally in agreement with the RNA-seq data (Fig 4C; Fig S4A). Notably, we detected strong expression of *Ln-CR*2 in E11.5 Ebf2-EGFP+ cells, and comparably lower or even no signals in Ebf2-EGFP- cells, indicating Ebf2-EGFP+ cell-specific expression (Fig 4C; Table S5). ISH on E11.5, E13.5 and E15.5 mouse embryonic brain sections revealed that *Ln-CR*2 was selectively expressed in the known CR neuron origins as Sp (Septum), CH (cortical hem) regions, as well as the TE (thalamus eminence), one of the extra-cortical origins of CR neurons, indicating cell-type specificity (Fig 4D). Interestingly, novel CR marker coding genes and lncRNAs identified from bulk RNA-seq results here were also enriched in single cell RNA-seq pseudo-temporal subpopulation C2, for instance, *Nhlh2*, *Ppp2r2c*, *Ln-CR4*, *Ln-CR7* and *Ln-CR9*. Some CR-specific genes like *B3gat1*, *Sncb* and CR-enriched lncRNAs as *2900079G21Rik* were found in C0346 (Fig 1E and F; Fig S1G; Table S2; Table S5). The consistency of bulk and single cell RNA-seq data sets allowed us to verify these novel genes’ CR neuron identities. On the other side, we also provided specific gene signatures and a range of putative novel markers for various cell states of CR neuron population.

**Figure 4.**
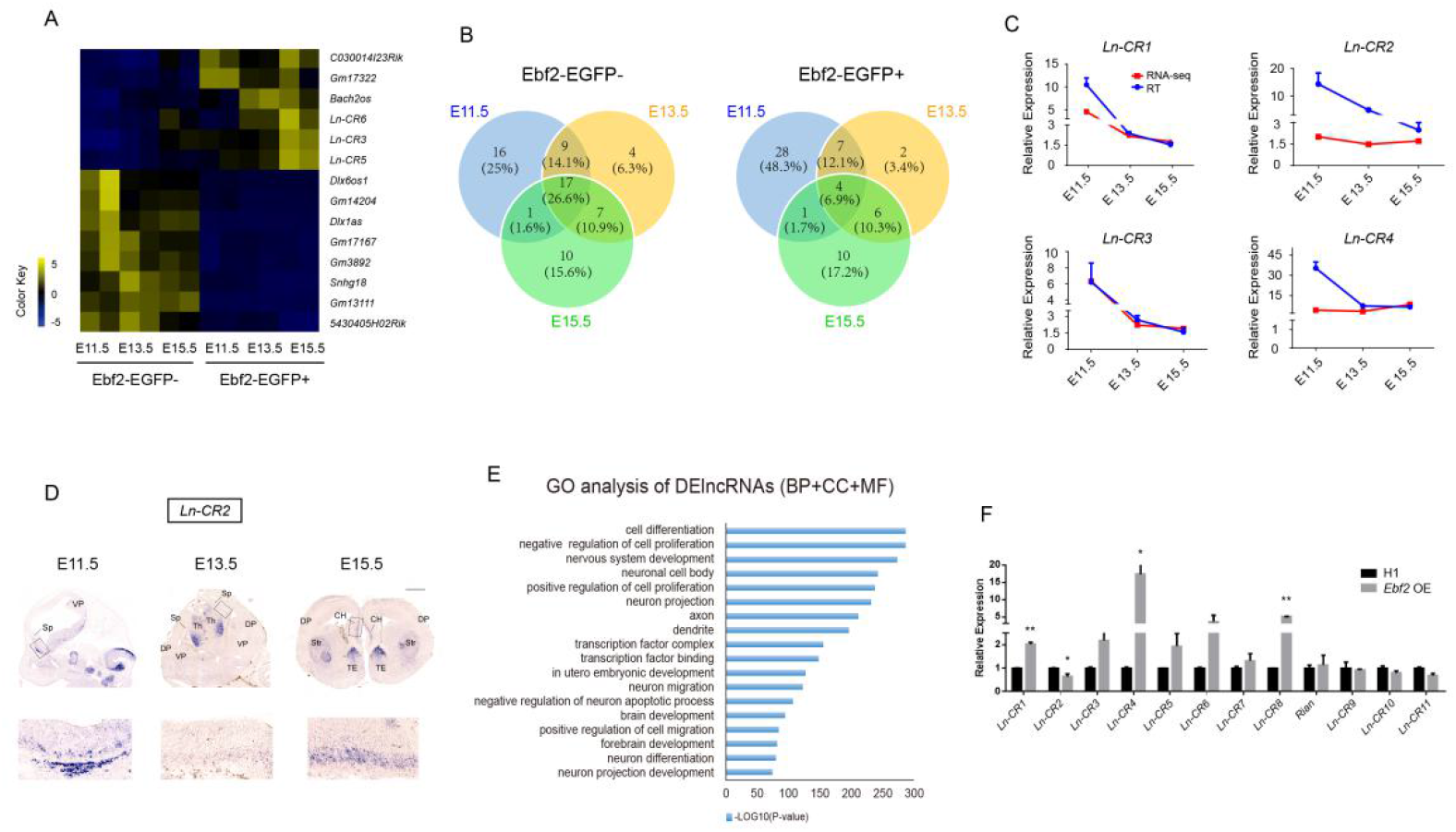
Characterization of differentially expressed lncRNAs in the Ebf2-EGFP+ and Ebf2-EGFP- cells. (A) Heatmap representing all differential expressed lncRNAs (FDR<0.05, p-value<0.05) in Ebf2-EGFP+ & Ebf2-EGFP-cell samples at the three embryonic stages. (B) Venn diagram displaying the extent of overlapping between DElncRNAs at the three embryonic stages. (C) The expression pattern of DElncRNAs between Ebf2-EGFP+ or Ebf2-EGFP- cells is consistent revealed by RNA-seq and qRT-PCR. (D) In situ hybridization showing the endogenous expression level of CR-specific lncRNA *Ln-CR*2 at E11.5, E13.5 and E15.5. Regions with specific expression are labeled and higher magnification images of boxed areas are shown below. Abbreviations: SP, Septum; VP, ventral pallium; Th, thalamus; DP, dorsal pallium; CH, cortical hem; Str, striatum; TE, thalamus eminence. (E) Enriched functional assessments (including biological processes, cellular components and molecular function) of highly expressed DElncRNAs in Ebf2-EGFP+ or Ebf2-EGFP-cell population (fold change ≥ 2). (F) qRT-PCR analysis comparing the expression levels of lncRNAs in NSC culture assay after virus infection with control (H1) or *Ebf2*-overexpression. Data represent mean ± SEM (n=4 independent experiments, *P<0.05, Student’s T test).

Furthermore, we performed GO analysis (for more details, see supplemental method) of DElncRNAs to predict their potential functions (Fig 4E). We found significant enrichments of neuronal development process-related GO terms, such as “nervous system development”, “neuron migration”, “negative regulation of neuron apoptotic process” and “neuron differentiation”, neuron identities-related GO terms such as “neuron projection”, “axon” and “dendrite”, also transcription factors regulation-related GO terms such as “transcription factor complex”, “transcription factor binding”. The functional annotation of DElncRNAs identified from our sequencing data suggests that these lncRNAs are potential players in the processes of neurogenic lineage progression, neuronal maturation and transcription factor regulation (Fig 4E; Table S6). Overexpression of *Ebf2* significantly up-regulated expression levels of some CR-specific DElncRNAs *Ln-CR1, Ln-CR4* and *Ln-CR8* but not others like *Ln-CR2* (Fig 1F; Fig 4F). We also performed knockdown of *Foxg1*, which significantly up-regulated *Ln-CR3* and *Ln-CR*5. Some lncRNAs like *Ln-CR*1 also had the trends of up regulation, while others had not, like *Ln-CR2* and *Ln-CR7*, implying that CR-specific lncRNAs might be involved in different regulatory pathways of CR neuron development (Fig S4B).

### Distinct H3K27ac and H3K27me3 histone modification patterns in Ebf2-EGFP+ cells

Histone modification is a critical determinant of transcriptionally active or silent chromatin states (57). To gain a comprehensive view of cell type-specific histone modification in early neural progenitor cells and differentiating neurons, we performed H3K27ac and H3K27me3 chromatin immunoprecipitation followed by high-throughput sequencing (ChIP-seq) in Ebf2-EGFP+ cells as early cortical development (E11.5, E13.5, E15.5) and Ebf2-EGFP- cells at E11.5 (mostly NSCs and neural progenitor cells at this stage). We analyzed the genome wide and gene-specific occupancy of these histone modifications in combination with the RNA-seq results on these cell types. First, our data shown strong concordance in H3K27ac and H3K27me3 bindings across two biological replicates (Fig S5A and B). According to a general strategy used to classify REs (regulatory elements) (58–60), we classified H3K27ac and H3K27me3 bound TSS (± 5kb) by the pattern of histone modification as TSS activation (monovalent H3K27ac), repression (monovalent H3K27me3) and poised (bivalent H3K27ac and H3K27me3). We observed bivalent H3K27ac and H3K27me3 bindings at 58.3% TSS sites of all detected genes in Ebf2-EGFP- cells (0.9% had only H3K27ac bindings and 4.5% had only H3K27me3 bindings). In contrast, the majority of genes in Ebf2-EGFP+ cells carried TSS activating modifications with strong H3K27ac and weak H3K27me3 enrichments (72.4% had only H3K27ac bindings, 0.5% had only H3K27me3 bindings, 16.4% had both) (Fig 5A and B). Such observations were consistent with previous studies that bivalent histone modification patterns formed a poised state to be ready for initiation of further developmental events in the ES cells or progenitor cells, while monovalent states presented in the differentiated cells indicated more organized states in the established specific cell identity (60, 61). Compared to E11.5 Ebf2-EGFP-progenitor cells, in which H3K27ac enrichments at TSS were the lowest while H3K27me3 bindings were the highest among the tested groups, H3K27ac enrichments at TSS in E11.5 Ebf2-EGFP+ cells were stronger and became increasingly higher while H3K27me3 bindings remain steadily at a lower level as development proceeds from E11.5 to E15.5 (Fig 5A and B). These patterns suggested that histone modification patterns were different between early differentiating neurons and neural progenitor cells, and dynamically changed as Ebf2-EGFP+ cells differentiated.

**Figure 5.**
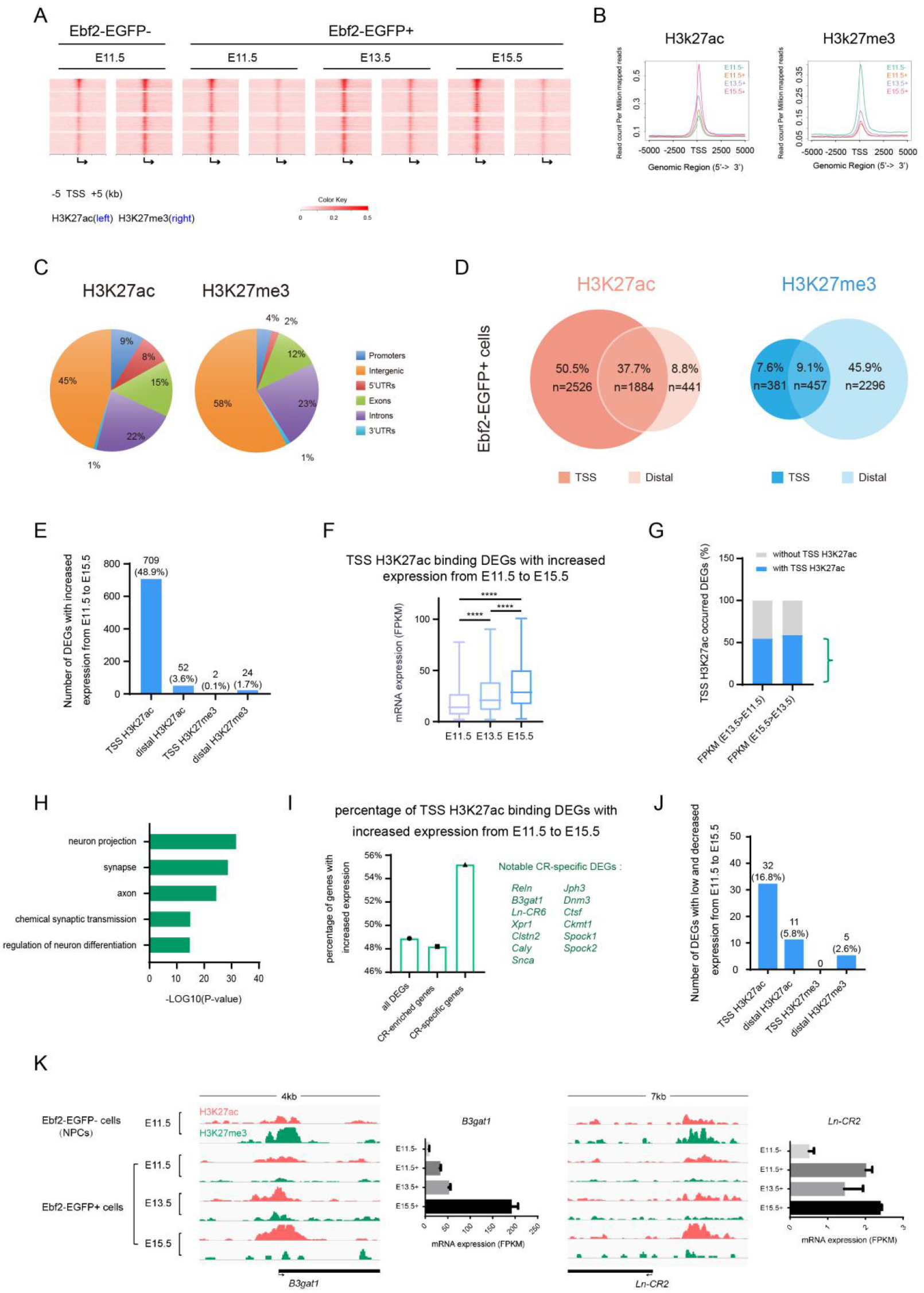
Genomic features of H3K27ac and H3K27me3 histone modification in the early differentiating neurons. (A) Heatmap representation of ChIP-seq signal density for H3K27ac and H3K27me3 for 10 kb centered on predicted Gene TSSs in Ebf2-EGFP-(E11.5) cells and Ebf2-EGFP+ (E11.5, E13.5 and E15.5). (B) A profile plot showing normalized H3K27ac (left) or H3K27me3 (right) enrichments around TSS of all detected genes in Ebf2-EGFP-(E11.5) cells and Ebf2-EGFP+ cells (E11.5, E13.5 and E15.5). (C) Genomic features of H3K27ac and H3K27me3-bound genes in Ebf2-EGFP+ cells. (D) Proportion of TSS and distal H3K27ac, H3K27me3 bindings distributed in DEGs in Ebf2-EGFP+ cells, respectively. (E) The number of DEGs that steadily increase expression level from E11.5 to E15.5 with TSS and distal region H3K27ac, H3K27me3 occupancies in Ebf2-EGFP+ cells. The percentage indicate the proportion of all E11.5 to E15.5 expression increased DEGs (with and without histone modification). Chi-square test showing the significant correlation between H3K27ac signals in the gene TSS regions and gene expression, P<0.01. (F) mRNA expression (FPKM) of DEGs with H3K27ac signals around TSS region increased from E11.5 to E15.5. Data represent mean ± SEM, N=334 genes, independent experiments, ****P<0.0001, T-test. (G) Percentage of DEGs that steadily increase expression level from E11.5 to E13.5, E13.5 to E15.5, with (blue) or without (grey) TSS H3K27ac enrichements, in the Ebf2-EGFP+ cell populations, that is, during CR neuron differentiation progression. (H) Functional GO analysis for those TSS H3K27ac binding DEGs in Ebf2-EGFP+ cells with steadily increase expression level from E11.5 to E15.5. Genes for analysis derived from Fig 5G. (I) Percentage of gene number (DEGs, CR-enriched DEGs and CR-specific DEGs) with positive correlation between expression (FPKM) and H3K27ac signals in the gene TSS region. Notable CR-specific genes are listed. (J) The number of DEGs that lowly expressed (FPKM<1) and steadily decrease expression level from E11.5 to E15.5 with TSS and distal region H3K27ac, H3K27me3 occupancies in Ebf2-EGFP+ cells. The percentage indicate the proportion of all E11.5 to E15.5 low and decreased expression DEGs (with and without histone modification). (K) The TSS regions of CR-specific genes *B3gat1*, and CR-specific lncRNAs *Ln-CR*2 demonstrate a similar pattern of histone modification during early embryonic development (bivalent in NPCs and become more activated in Ebf2-EGFP+ cells from E11.5 to E15.5).

We further analyzed the genomic location of H3K27ac and H3K27me3 bindings in Ebf2-EGFP+ cells (Fig 5C; Fig S5C-H). We found that the majority of H3K27ac and H3K27me3 binding peaks were located in the intergenic and intron regions, while less than 10% of peaks distributed in the promoter regions (Fig 5C; Fig S5C and D). At E11.5, the genomic distribution patterns of H3K27ac and H3K27me3 in Ebf2-EGFP- cells were similar to that in Ebf2-EGFP+ cells (Fig S5E-H).

Interestingly, a distinct histone modification pattern emerged when we focused on the TSS regions (within ±5kb) and distal regions (within 100kb) of DEGs. TSS and distal regions were reported important regulatory potentials in the development system (62, 63), so we concentrated on these two regions of DEGs in Ebf2-EGFP+ cells. We found that for the H3K27ac bindings, 88.2% (4410) genes in Ebf2-EGFP+ cells had peaks around TSS, while 46.5% (2325) had peaks around distal region. Interestingly, the majority of genes either had H3K27ac peaks only on TSS (50.5%, 2526 genes) or on both TSS and distal region (37.7%, 1884 genes), while 8.8% (441) had enrichments only on distal region (Fig 5D). Conversely, more than half (55.0%, 2753) genes had H3K27me3 peaks on distal regions, 16.7% (838) on TSS, 9.1% (457) on both TSS and distal regions, 7.6% (381) genes had bindings only on TSS (Fig 5D), showing that more genes occupied with H3K27me3 enrichments on distal region than on TSS, which is quite different from H3K27ac pattern. Hence, DEGs might be activated in the Ebf2-EGFP+ cells as more genes were occupied with H3K27ac enrichments, especially some CR marker genes as *Calb2*, *Trp73*, *Tbr1*, and novel CR-specific coding genes and lncRNAs identified in this study, for example, *Sncb*, *Ppp2r2c*, *Jph3*, *Ln-CR2*, *Ln-CR3*. The histone modification pattern as DEGs with H3K27ac at TSS region and H3K27me3 at distal region in Ebf2-EGFP+ cells might contribute to maintain and promote CR neuron differentiation.

### Differentiation genes are facilitated during CR neuron specification accompanied by TSS H3K27ac bindings

It has been shown that chromatin state around TSS or distal region is a strong indicator of RE function (60, 62, 63). We noticed that among 1555 CR-enriched genes, 97.1% of them were occupied by H3K27ac bindings (Fig S5I). Distal H3K27ac bindings are known to activate gene expression in many development systems including in neural system (62, 63). Interestingly, we found that 46.5% (2325) of DEGs had distal H3K27ac bindings in Ebf2-EGFP+ cells (Fig 5D), however only 52 genes showed increased expression level from E11.5 to E15.5 (Fig 5E and F). In contrast, we found that genes with H3K27ac signals surrounding TSS regions tended to have increasing expression as development proceeded. Genomically, 88.2% (4410) DEGs carried TSS H3K27ac bindings in Ebf2-EGFP+ cells (Fig 5D), 45.7% (2288) DEGs had consistent TSS H3K27ac modification across the three stages, and 31.0% (709) of them showed stably increased expression from E11.5 to E15.5 (48.9% of all increased DEGs), suggesting H3K27ac modification around TSS significantly increased temporal genes expression (Chi-squared test: p-value < 0.01) (Fig 5E and F). These genes significantly encoded early differentiating neuron characters and functions, like “neuron projection”, “synapse”, “axon”, “chemical synaptic transmission”, “regulation of neuron differentiation” (Fig 5G and H). Genes accompanied by TSS H3K27ac bindings across three developmental time points also occupied 75% (93) of highly expressed C2 genes (top 20%) from single cell pseudotime lineage (Fig S5J), implicating their constraints to CR neuron termination. For CR-enriched DEGs, 340 with TSS H3K27ac surroundings showed steadily increased expression along the time period (48.2% of all increased CR-enriched DEGs). For CR-specific DEGs, 32 were such genes (55.2% of all increased CR-specific DEGs), including *Reln*, *B3gat1*, *Ln-CR6*, *Xpr1*, *Clstn2*, *Caly*, *Snca*, *Jph3*, *Dnm3*, *Ctsf*, *Ckmt1, Spock1, Spock2* (Fig 5I). We wondered whether TSS H3K27ac also contributed to repress gene expression, however, only 1.4% (32) of consistent TSS H3K27ac binding DEGs showed low (FPKM<1) and decreased expression from E11.5 to E15.5 (16.8% of all low and increased expressed DEGs) (Fig 5J). Hence, promoter but not distal region H3K27ac modification contribute to up-regulate gene expression, especially CR-specific genes along differentiation. Besides, we noticed that 16.7% (838) and 55% (2753) DEGs had TSS and distal H3K27me3 bindings in Ebf2-EGFP+ cells, respectively (Fig 5D). However, we found only 2 and 24 genes with increased expression at the three stages (Fig 5E). On the other hand, we also found distal H3K27me3 did not suppress gene expression because only 11 low expressed genes under their modification (Fig S5K). Therefore, the data shown that, unexpectedly, H3K27ac activation at promoter but not distal region in Ebf2-EGFP+ cells facilitated early neuron differentiation. Here are examples reflecting H3K27ac and H3K27me3 modifications involvements in CR-specific coding genes and lncRNAs (e.g. *B3gat1*, *Ln-CR2, Sncb, ln-CR5*) differential expression (Fig 5K; Fig S5L). Their expression increase in Ebf2-EGFP+ cells from E11.5 to E15.5, accompanied by increasingly stronger H3K27ac enrichments at TSS regions during these stages. Meanwhile, these genes are expressed at low levels in Ebf2-EGFP- cells on E11.5, corresponding to H3K27me3 enrichments at TSS of these genes in Ebf2-EGFP- cells (Fig 5K; Fig S5L). *Ln-CR*2, for example, which is increased in Ebf2-EGFP+ cells from E13.5 to E15.5, exhibits a change in histone modification with low H3K27ac peaks on the TSS region on E13.5 but high H3K27ac peaks on E11.5 and E15.5 (Fig 5K).

### Functional validation of CR-specific lncRNAs from *In Vitro* and *In Vivo*

In this study we have identified CR-specific lncRNAs, however the functional roles of these lncRNAs in neurogenesis have not been studied before. Using lentivirus-mediated shRNA silencing and overexpression, we performed *in vitro* and *in vivo* functional assays. We cultured single E11.5 cortical progenitor cells in adherent conditions for 5 days (DIV5), then fixed the culture and stained for neuronal markers β-III tubulin (immature neurons) and *Reln* (CR neurons) (Fig 6A; Fig S6A). In the control culture 29.89% of the cells were β-III tubulin+ neurons, of which 3.16% were Reln+ (Fig 6B; Fig S6B). Overexpression of *Ln-CR*1, *Ln-CR*2 and *Ln-CR*3 increased β-III tubulin+ cells and reduced total cell number (Fig 6C and D; Fig S6C and D). Notably, Reln+ neurons were increased to 11.56% by *Ln-CR*1 overexpression, 6.09% and 6.83% by *Ln-CR*2 and *Ln-CR*3 overexpression, respectively (Fig 6B; Fig S6B). Interestingly, the number of small clones with less than 4 cells and all β-III tubulin+ increased in CAG-*Ln-CR*1, CAG-*Ln-CR*2 and CAG-*Ln-CR*3 compared to CAG-control, while the number of clones with more than 8 cells was much fewer than that in control (Fig 6C; Fig 6E; Fig S6C; Fig S6E). These results suggest that overexpression of CR-specific lncRNAs promotes neuronal differentiation. Moreover, we also observed that overexpression of *Ln-CR*1 altered the morphology of Reln+ neurons, increased whole processes length, branch depth and branch levels, while no significant differences were detected in neuron soma size (Fig 6F; Fig S6F). During the postnatal stage, CR neurons develop more complex morphology (30). We further analyzed the effect of *Ln-CR*1 on cell morphology in an *ex-vivo* culture system using the wholemount cortical hemispheres isolated from E15.5 Ebf2-EGFP+ embryos and cultured for 10 days (DIV10) to mimic late embryonic to postnatal development. Compared to control, we found that overexpression of *Ln-CR*1 increased the morphological complexity of CR neurons, leading to more filopodia, spine-like structures and bouton-like structures (Fig S6G and H). We observed similar results in P3.5 embryos that *Ln-CR*1 overexpressed by in utero electroporation into the layer 1 of the E15.5 cerebral cortex (Fig 6H).

**Figure 6.**
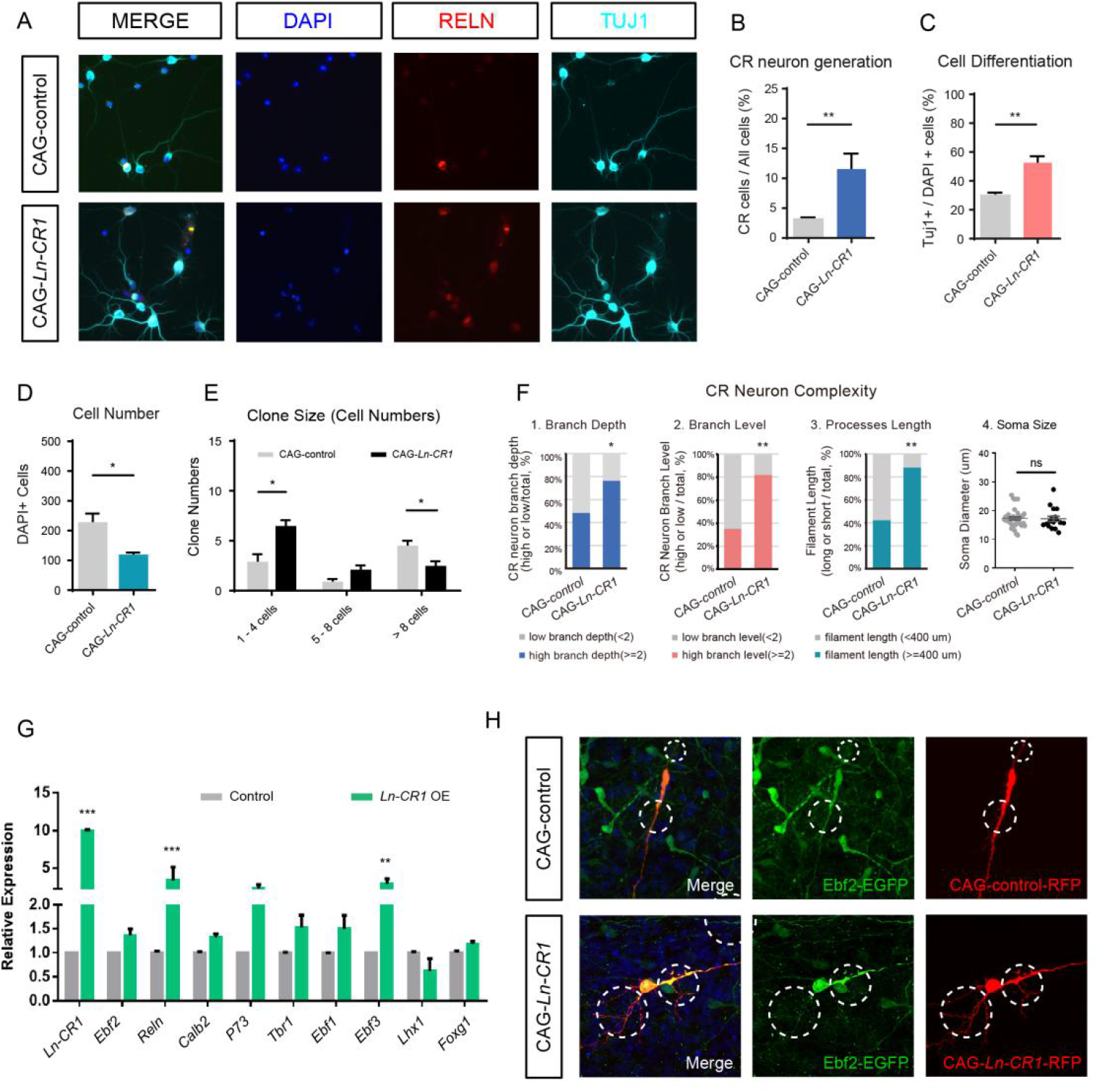
Functional validation of CR-specific lncRNAs *Ln-CR*1 *In Vitro* and *In Vivo.* (A) Immunostaining of NSCs treated with control (CAG-control) or overexpression lentiviruses for CR-specific lncRNA *Ln-CR*1 (CAG-*Ln-CR*1) after in vitro culture for 5 days. RELN (red), TUJ1 (cyan), or DAPI (blue) for CR neurons, neurons, and nucleus, respectively. Scale bar, left 3 rows 150 µm, right 50 µm. (B-E) Overexpression of CR-specific lncRNA *Ln-CR*1 significantly increased CR neuron number, promoted NSC differentiation and generated more neuron-like small cell clones. All the experiments were performed 4 times and 8 randomly selected microscopy fields were counted each time. Data represented mean ± SEM. p values indicated were calculated by Student’s t test (unpaired). **p<0.01, *p<0.05. (F) Quantification of process branch depth, branch levels and total dendrites length and soma size of CR neurons after overexpression of CR-specific lncRNA *Ln-CR*1 compared to control. All the experiments were performed 3 times and 8 randomly selected microscopy fields were counted each time. Data represented mean ± SEM. p values indicated were calculated by Student’s t test (unpaired). **p<0.01, *p<0.05. (G) Quantification of CR neuron molecular markers as *Reln*, *Ebf2*, and *Calb2* at 72h after lentiviral transduction in the NSC culture assay comparing the effect of overexpression of *Ln-CR*1 to control. Data represented mean ± SEM (**p<0.01, *p<0.05, n>3, Student’s T test). (H) Comparison of dendritic spine, filopodia and processes in cortical wholemounts overexpressing *Ln-CR*1 (CAG-*Ln-CR*1) to CAG-control. Scale bar, 20 µm. Ebf2-EGFP+ cortical wholemounts were collected from P3 pups after *in utero* electroporation to cortex layer 1 on E15.5.

To uncover how overexpression of these CR-specific lncRNAs molecularly affected CR neuron development, we assessed the mRNA levels of several known CR neuron-related genes by qPCR (Fig 6G; Fig S6I). We found genes highly expressed in CR neurons such as *Reln*, *Ebf2*, *Calb2* and *Trp73*, *Ebf1*, *Ebf3*, *Tbr1*, *Nhlh2* and *Lhx1*, were selectively altered when overexpressing CR-specific lncRNAs candidates *Ln-CR*1, *Ln-CR*2 and *Ln-CR*3. While typical CR markers represent major CR neuron populations as *Reln*, *Ebf3*, *Ebf2* and *Trp73* were significantly upregulated under one specific lncRNA overexpression(Fig 6G; Fig S6I), implying that these lncRNAs might have great potentials in regulating the molecular pathways for CR neuron differentiation.

## Discussion

In this study, we characterized the dynamic transcriptomic profile of early neurogenesis and identified gene expression signatures associated with preplate neuronal differentiation and subsequent establishment of CR cell properties. We also demonstrated that E15.5 CR cells could be classified into eight subpopulations with three distinct maturation neuron states by single cell RNA-seq analysis, which has not been reported before. Furthermore, we provided the first neuron subtype-specific profile of histone modifications, revealing active or repressive histone modification states in the early differentiating neurons that were correlated with gene expression level. We have identified novel lncRNAs expressed in CR cells and some of them were functionally important for CR cell differentiation. Our findings provide a comprehensive molecular depiction of the early neuronal differentiation events, shedding new light on the mechanism underlying preplate formation and differentiation of CR cells, the first step of cortical neurogenesis.

### Molecular transition from preplate to CR neurons

As cortical neurogenesis proceeds, the preplate is physically separated into the subplate and the layer 1 by migrating cortical plate neurons. We first identified over 1044 coding genes and 40 lncRNAs with higher expression in the newly differentiated preplate neurons (Ebf2-EGFP+) at E11.5 (Fig 2D; Fig 4B; Table S2; Table S5). At E13.5, 370 genes continue to show enrichments in preplate neurons, while additional 20 genes also have higher expression in subplate neurons. A subset of these genes continue to be differentially expressed in the Ebf2-EGFP+ cells at E15.5, while the total DEGs at E15.5 are much less than earlier stages, suggesting further specification of the CR cell fate. Indeed, we found fewer overlaps between the confirmed subplate neuron-specific genes at this age (18) (28 genes overlap, 2.3% of total DEGs at E15.5), confirming the molecular segregation of preplate neurons into the subplate and layer 1 by E15.5. The fact that the number of DEGs is the highest at E11.5 reflects the degree of the difference between the two populations: At E11.5, all Ebf2-EGFP+ cells are early differentiating neurons, while the Ebf2-EGFP- cells are mostly cortical progenitor cells. As the Ebf2-negative population includes both cortical plate neurons and neural progenitor cells at E13.5 and E15.5, genes common to neurons might not have significant changes.

This dynamic transcriptional profile may underlie the progression of preplate differentiation to the acquisition of unique molecular identity of CR neurons and subplate. For example, 166 genes are enriched in E11.5 and highly expressed in preplate, then later become restricted in CR neurons, while 851 genes which are also enriched in E11.5 but no longer specifically expressed in layer 1 at E15.5.

Our transcriptome analysis also reveals a surprising enrichment of genes categorized in the GO terms (Fig 3B-D) such as “neuron projection”, “neuron differentiation”, “axonogenesis”, even “synapse” and “synaptic transmission” as early as E11.5. Terms such as “gate channel activity” and “ion channel activity” also appear before E15.5. Interestingly, whole-cell patch-clamp recording showed an inward sodium current in putative CR cells as early as E12 (64). In addition, dynamic expression of K+ channels and Ca2+ channels were also described in pre- and postnatal CR cells (12, 64–66). We found that Ebf2-EGFP-positive cells at E11.5 already expressed neurotransmitter receptors including ionotropic and metabotropic glutamate receptors, AMPA glutamate receptors, at E11.5 (Table S2, Table S3). Interestingly, it has been reported that at very early stages of neural development, subplate neurons form transient synapses with axons coming from the subcortical regions to guide later formation of cortico-subcortical connectivity (67, 68). Our results suggest that populations of early differentiating neurons, including Cajal-Retzius cells, are equipped with machineries suited for electric or chemical synaptic connections shortly after they are born. It’s possible that functional connections are established earlier than currently recognized. Alternatively, expression of these synapse-related proteins may not necessarily confer electrophysiological properties, but rather prime early differentiating neurons to be ready for synaptic connection once they encounter with other cortical plate neurons.

Disease-related genes are highly expressed during early differentiation period (18, 69). Abnormality in the process of preplate formation and splitting are associated with neurological disorders including mental retardation, epilepsy, and autism (8). Interestingly, schizophrenia has a highly susceptible stage between 6-8 weeks gestation, the time when preplate differentiation occurs (70). These findings suggest the importance of proper regulation during the early differentiation process.

### Single-cell RNA-seq analysis reveals distinct developmental states within pure CR neuron populations

Pseudo-time analysis of single cell RNA-seq data has offered a way to further dissect the transitional difference in gene expression within a heterogeneous population. Here, we reported that by pseudo-time analysis, CR neurons could be classified into 3 subgroups. The first group consisted of activated cluster 1, 5 and 7 genes, which were highly expressed at the beginning of pseudo-temporal axis. We found that genes in this group are associated with characteristics of cells located in the upper VZ and lower cortical plate such as *NeuroD1*, representing transitioning from neural progenitor cells to newborn neurons (34, 35, 71). The second group is enriched with cluster 0, 3, 4, 6 genes that were activated during the middle of pseudotime trajectory. These genes were categorized under “neural differentiation” “neurite outgrowth” and “migration” terms using DAVID analysis. The third group along the pseudo-time line consisted of mainly cluster 2 genes, which includes many CR marker genes, synapse and axonogenesis-related genes. Interestingly, we found that genes first expressed in neural progenitor cells then in cortical plate neurons, such as *Neurod1* and *Rnd2*, were expressed higher first, then down-regulated in the third group, suggesting when cells undergo fate decisions to become CR neurons, genes related to other neuronal identity would need to be suppressed at the later differentiation stage. Indeed, transcription factors, such as EBF2 and ZIC2, expression levels of which increase as CR neurons differentiate, and FEZF2, which is highly expressed in the intermediate state, could form dynamic regulatory cascades via corporately repressing non-CR genes and inducing CR-specific genes to promote the fine molecular transition from a new-born neuron to a fully committed subtype. Further studies will help resolve the biological relevance of these analysis-acquired developmental trajectory of gene expression. Overall, our study provides a comprehensive profile of gene expression for the early born neurons in the cerebral cortex at population and single cell level, unraveling the molecular dynamics during early neurogenesis.

## Materials and Methods

### Mice

All animal protocols used in this study were approved by the IACUC (Institutional Animal Care and Use Committee) of Tsinghua University and performed in accordance with guidelines of the IACUC. The laboratory animal facility has been accredited by AAALAC (Association for Assessment and Accreditation of Laboratory Animal Care International). For timed mice mating, the day of the vaginal plug was considered E0.5.

### Mapping and transcripts assembling

Sequencing reads were mapped to mouse genome using bowtie with parameter “-v 2 -X 350 -fr -m 10 -S -k 1”. After mapping sequencing reads to the mouse genome, we assembled the transcripts using Cufflinks (72). We assembled transcripts into known gene models or novel gene models guided by known gene annotations. Known gene annotations were downloaded from Ensembl (Mus_musculus.GRCm38.74). The assembled transcripts were then merged by the Cuffmerge utility provided by the Cufflinks package. Then the merged transcripts were concurrently annotated by Cuffcompare program in Cufflinks suite, of which known transcripts were used as reference to screen for novel lncRNAs. Only transcripts with tag “intergenic” and “intronic” were retained. Transcripts shorter than 200nt or with FPKM lower than 0.3 were filtered out from the transcripts set.

### Tissue collection and RNA In Situ hybridization

Mouse brains at various developmental stages were collected and fixed in 4% paraformaldehyde (PFA) at 4°C for 4h. After briefly rinsing in PBS, brains were dehydrated in 30% sucrose in PBS at 4°C until the time they sank, and embedded in OCT (Sakura Finetek, Torrance, CA). Frozen brains were cryosectioned to 16 μm and mounted to coverslip. In situ hybridization was performed according to instructions as previously described (Wigard et al., 2006; Gregor et al., 2007; Robert et al., 2007). Digoxigenin-labeled RNA probes were designed based on lncRNA sequences determined by Cufflinks and Ensembl. Images were taken on a Zeiss Slide Scanner (Axio Scan. Z1) fluorescent microscope.

### Chromatin-immunoprecipitation and sequencing (ChIP-Seq)

ChIP assays were performed using mice primary Ebf2-EGFP positive cells at E11.5, E13.5, E15.5 and Ebf2-EGFP negative cells at E11.5, with antibody against H3K27ac (Ab4729, Abcam) and H3K27me3 (Ab6002, Abcam), experimental procedures were as previously described (73). Briefly, Ebf2-EGFP positive and negative cells (∼1-5 million cells) were sorted and collected from FACS (BD Influx^TM^), and fixed by 0.4% (v/v) formaldehyde at room temperature (RT) for 10 min. The reaction was stopped by adding 0.125 M glycine for 5 min. After rinsing cells three times with cold PBS, the nuclei were resuspended in Nuclei Lysis Buffer (50 mM Tri-Cl, pH8.1, 10 mM EDTA, 1% SDS, 1 mM DTT, 1 mM PMSF and Sigma protease inhibitor cocktail, RT) and incubated on ice for 10 min. Nuclear suspension were sonicated 6” on, 15” off, 20 cycles (power set at 25%) on ice. The fragmented chromatin was diluted for 5 folds in ChIP Dilution Buffer (16.7 mM Tri-Cl, pH8.1, 167 mM NaCl, 1.2 mM EDTA, 1.1% Triton X-100, 1 mM PMSF, and Sigma protease inhibitor cocktail). Then 5 ug IP antibody was added and incubated overnight at 4 °C. Chromatin-protein complex was pulled down with Dynabeads protein G (Cat#10004D, Invitrogen) for 2h at 4°C with rotation and followed by 5 sequential washes as the following: 1. low salt wash buffer (20 mM Tri-Cl, pH8.1, 150 mM NaCl, 1 mM EDTA, 1% Triton X-100, 0.1% SDS, 0.1% Na-deoxycholate); 2. high salt wash buffer (20 mM Tri-Cl, pH8.1, 500 mM NaCl, 1 mM EDTA, 1% Triton X-100, 0.1% SDS, 0.1% Na-deoxycholate); 3. LiCl wash buffer (10 mM Tri-Cl, pH8.1, 250 mM LiCl, 1 mM EDTA, 0.5% NP-40, 0.5% Deoxycholic acid (Na salt)); and the last two washes with TE buffer (10 mM Tri-Cl, pH8.1, 1 mM EDTA). Immunoprecipitated complex was eluted from the antibody in elution buffer (1% SDS, 50 mM TrisCl (pH 8.1), 1 mM EDTA) by votexing once at 65°C for 20min and once at RT. Eluted chromatins were incubated at 65°C overnight for reverse crosslinks and chromatin fragmented purification with PCR column purification. ChIP-Seq libraries were prepared using NEB Next Ultra^TM^ Ⅱ ChIP-Seq Library Prep Kit (Cat#E7645S & Cat#7335S, NEB) according to the manufacturer’s instructions. ChIP-seq data were aligned to mouse genome mm10, performing Bowtie v. 1.1.2 on the Galaxy platform with the following options: -v 3 –M 1 −5 15. Resulting output SAM files were converted to BAM format, and finally converted to BED files. Peaks were called using MACS (Model-Based Analysis of ChIP-Seq), with a p-value cutting off of 1e-5.

### Data Availability

The RNA-seq, ChIP-seq and scRNA-seq data in this study are under submission to the Sequence Read Archive (SRA; https://www.ncbi.nlm.nih.gov/sra) under submission number SUB4371065.

## Acknowledgments

We thank Dr. Zhengang Yang and Dr. Chunyang Wang at Fudan University Collaborative Innovation Center for Brain Science for their assistance of In Situ hybridization experiments in this study. We thank Yuchuan Wang, Chao Di and Xinqiang Ding at Tsinghua University for their help of bioinformatic analysis. We thank for all members of Shen lab for their helpful discussion during the preparation of manuscript. We are also grateful to Center of Biomedical Analysis at Tsinghua University.

